# Ca^2+^-induced mitochondrial ROS regulate the early embryonic cell cycle

**DOI:** 10.1101/223123

**Authors:** Yue Han, Shoko Ishibashi, Javier Iglesias-Gonzalez, Yaoyao Chen, Nick R. Love, Enrique Amaya

**Author notes:** Current address: STEMCELL Technologies UK ltd, Building 7100, Cambridge Research Park, Beach Drive, Waterbeach, Cambridge, UK, CB25 9TL. Current address: Stanford University School of Medicine, Stanford, CA 94305, USA. Co-first author (Yue Han, Shoko Ishibashi).

## Abstract

While it has long been appreciated that reactive oxygen species (ROS) can act as second messengers in both homeostastic and stress response signaling pathways, potential roles for ROS during early vertebrate development have remained largely unexplored. Here we show that fertilization in *Xenopus* embryos triggers a rapid increase in ROS levels, which oscillates with each cell division. Furthermore, we show that the fertilization induced Ca^2+^ wave is both necessary and sufficient to induce ROS production in activated or fertilized eggs. Using chemical inhibitors, we identified mitochondria as the major source of fertilization induced ROS production. Inhibition of mitochondrial ROS production in early embryos results in cell cycle arrest, in part, via ROS dependent regulation of Cdc25C activity. This study reveals for the first time, a role for oscillating ROS levels in the regulation of the early cell cycle in *Xenopus* embryos.

**Highlights:**

- ROS, including hydrogen peroxide, are produced after fertilization in *Xenopus*
- Ca^2+^ signaling after fertilization induces ROS production in mitochondria
- Mitochondria are the major source of oscillating ROS levels
- ROS regulate Cdc25C activity and the early cell cycle

## INTRODUCTION

Reactive oxygen species (ROS), which include the superoxide anion (O_2_^•−^), hydroxyl radical (OH^•−^), hydrogen peroxide (H_2_O_2_) and singlet oxygen ^1^O_2_), have various functions/activities in cells and tissues. ROS have several sources in the cell, including monoamine oxidases, xanthine oxidase, NADPH oxidases (NOXes) and several mitochondrial complexes, which are part of the electron transport chain (ETC) (Bedard and Krause, 2007; Cantu-Medellin and Kelley, 2013; Kaludercic et al., 2014). In addition mechanisms exist to change some forms of ROS into others. For example, superoxide dismutase (SOD) uses O_2_^•−^ as a substrate to generate H_2_O_2_, which is a more stable, diffusible and less harmful ROS, and can act as a second messenger in the cell (Burgoyne et al., 2013). Once generated, H_2_O_2_ has the capacity to modulate the activity of protein phosphatases via oxidizing catalytic cysteine residues, which can in turn enhance kinase-driven signaling pathways, such as MAP kinase pathways (Östman et al., 2011).

Embryonic development is triggered by sperm entry during fertilization, which activates the egg by generating a Ca^2+^ wave and increasing oxygen uptake and ATP synthesis. This increase in oxygen consumption was first noted over 100 years ago in sea urchin eggs (Warburg, 1908) and was later found to correlate with increased ROS production (Foerder et al., 1978). More recently, it was found that the dual oxidase protein (Udx1) is responsible for the generation of ROS following fertilization in sea urchins zygotes (Wong et al., 2004). Notably Wong and colleagues also showed that Udx1 is present throughout early development and that inhibition of ROS generation using diphenyleneiodonium (DPI) had the capacity to block cell division (Wong and Wessel, 2005). This suggested that ROS may play an important role in the control of the early embryonic cell cycle, but the mechanisms by which ROS regulate the cell cycle and, moreover, whether ROS play similarly essential roles during early vertebrate embryonic development remains unknown.

In early *Xenopus* embryos, the cell cycle is driven by an autonomous oscillator which is cytokinesis independent (Hara et al., 1980; Murray and Kirschner, 1989). The cyclin B/cyclin-dependent kinase 1 (Cdk1) complex is a master regulator for entry into mitosis. Accumulating Cyclin B levels activate Cdk1, which in turn activates Cdc25C phosphatase, which then dephosphorylates the inhibitory phosphorylated Thr 14 and Tyr 15 in Cdk1, resulting in activation of cyclin B/Cdk1 complex. This positive feedback loop ensures entry into mitosis. Conversely, Cdk1 also generates a negative feedback loop by activating the anaphase-promoting complex or cyclosome (APC/C) and the E3 ubiquitin ligase APC/C with co-activator, Cdc20 that promotes degradation of cyclin B, thus ensuring the exit of mitosis. These positive and negative feedback loops are thought to constitute an ultrasensitive bistable circuit to generate a cell cycle oscillator (Ferrell, 2013).

Mitochondria are important organelles that generate ATP in aerobic eukaryotes and participate in other aspects of cellular metabolism and cell signaling. It has been thought that mitochondria produce ROS as a by-product, however recent studies have shown that mitochondrial ROS (mtROS) can mediate intracellular signaling. For instance, mtROS generated in complex III was shown to be essential in antigen-specific T cell activation *in vivo* (Sena and Chandel, 2012). In fact, there are at least 11 sites in mitochondria, which produce ROS (Brand, 2016; Mailloux, 2015). Although mitochondrial Complex I and III are thought to be the major sources of mtROS, their contributions to overall ROS production appear to differ between species, organs, tissues and mitochondrial subpopulations. For example, Complex III produces most of the ROS generated by heart and lung mitochondria, while Complex I is responsible for most of the ROS produced in brain mitochondria *in vitro* (Barja and Herrero, 1998; Turrens and Boveris, 1980; Turrens et al., 1982). How or whether mitochondrial ROS-producing enzymes affect cellular embryonic processes *in vivo*, however, has thus remained largely unknown.

Using a transgenic Xenopus line expressing a H_2_O_2_ indicator, HyPer, we found that fertilization induces a rapid increase in ROS levels *in vivo*. Using this assay, we then explore both the molecular mechanisms that trigger the burst of ROS after fertilization and how these ROS effect early embryonic development. We show that Ca^2+^ induces ROS production after fertilization and that the major source of ROS during the early embryonic development are the mitochondria. We then show that mitochondrial ROS production oscillates with the cell cycle, and this oscillation participates in the regulation of the cell cycle in early Xenopus embryos, at least partly through inactivation of the cell cycle phosphatase, Cdc25C.

## RESULTS

### Fertilization induces increased ROS levels in *Xenopus* oocytes

We previously showed that *Xenopus* tadpole tail amputation induces sustained ROS production, which is necessary for successful tail regeneration (Love et al., 2013). For that study, we generated a transgenic *Xenopus laevis* line that ubiquitously expressed the H_2_O_2_ sensor HyPer (Belousov et al., 2006; Love et al., 2013; 2011). Serendipitously, when we examined the eggs from the transgenic females of this transgenic line, we found that HyPer was expressed maternally. We subsequently filmed fertilization using these HyPer expressing eggs and found that fertilization induced an 85% increased HyPer ratio (n = 11, 1-cell compared to egg, p = 0.001, Wilcoxon matched-pairs signed rank test), indicating an increased production of ROS that was sustained throughout early development (Figures 1A and 1B; Movie S1 available online). To examine the mechanisms that regulate the fertilization-induced ROS production in *Xenopus* embryos oocytes, we injected non-transgenic, immature oocytes with HyPer mRNAs. We then allowed these mRNAs to translate and induced maturation in a subset of these injected oocytes by treating them with progesterone (Figure 1C). While HyPer expressing immature oocytes failed to activate following pricking nor did they show a change their ROS levels (n = 33, p = 0.2, 20 min compared to 0 min, paired t-test) (Figures 1D and 1E), HyPer expressing mature oocytes were immediately activated by pricking, and these activated mature oocytes showed a dramatic increase in ROS levels (43.5 % increase in ratio, n = 28, p < 0.0001, 20 min compared to 0 min, paired t-test), mimicking the increase ROS levels we had seen following fertilization (Figures 1F and 1G). We also found that prick activation of unfertilized eggs obtained from transgenic HyPer-expressing females also resulted in dramatic increase in ROS production (n = 60, p < 0.0001, 20 min compared to 0 min, paired t-test) (Figures S1A and S1B). Since HyPer has also been shown to be pH sensitive, oocytes were also injected with SypHer RNA, which encodes a pH sensitive, but ROS insensitive mutant version of HyPer (Poburko et al., 2011). While we detected a slight increase of SypHer ratio in prick activated eggs (3.5 % increase in ratio, n = 27, p < 0.0001, 20 min compared to 0 min, Wilcoxon matched-pairs signed rank test) (Figures 1H and 1I), corresponding to a pH increase from 7.37 to 7.42, consistent with a previous finding (Webb and Nuccitelli, 1981), the change was considerably less than that seen with the ROS sensitive HyPer version. Therefore we concluded that the change of HyPer ratio following fertilization is primarily due to an increase in ROS levels, rather than a change in pH. Thus both fertilization or prick activation of mature oocytes and eggs led to a significant increase in ROS production in the activated eggs/zygotes.

**Figure 1.**
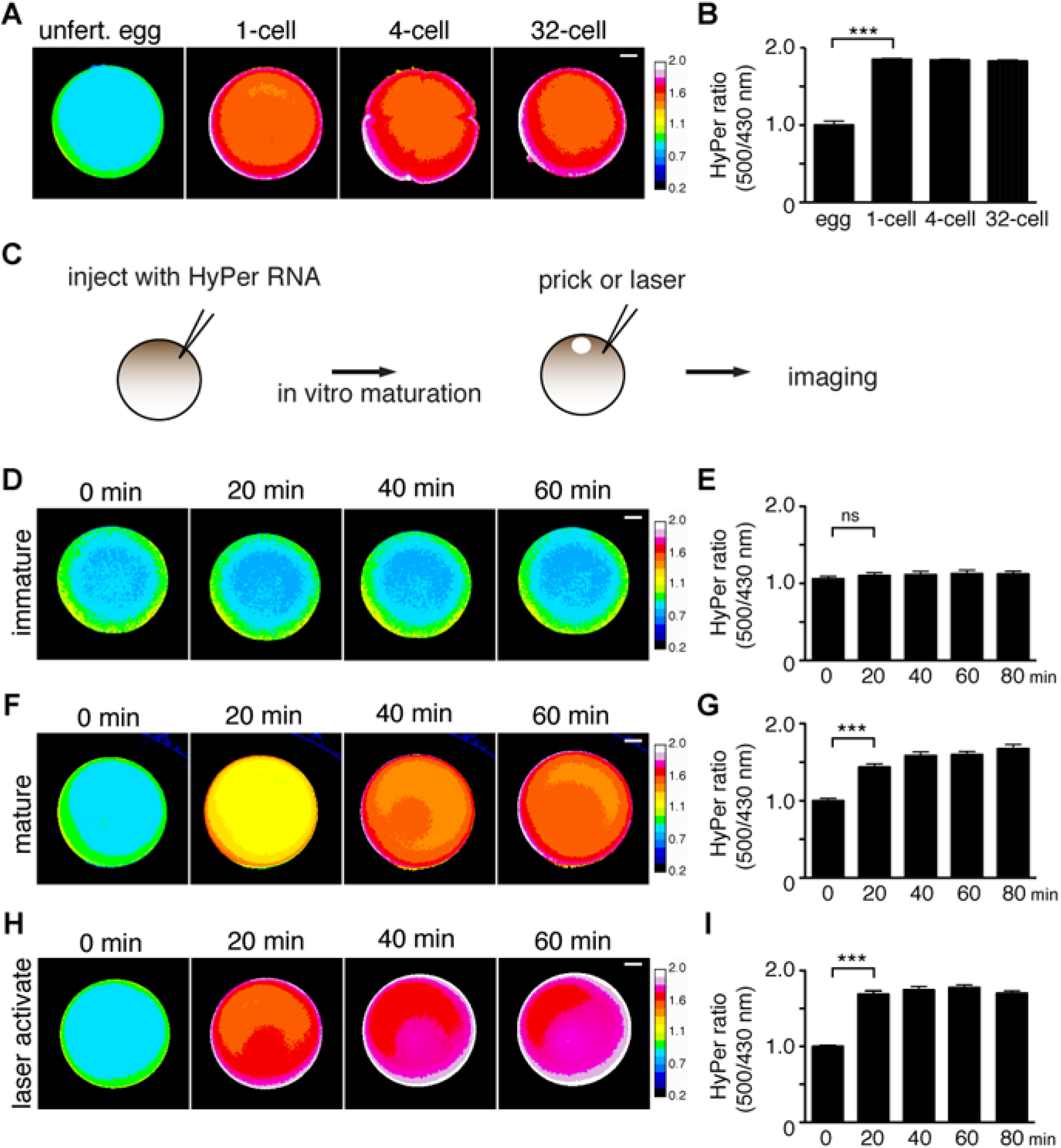
Fertilization and injury triggers a substantial increase in ROS levels. (A) HyPer ratio images (500/430 nm) showing a ROS production in transgenic embryos expressing HyPer. See also Movie S1. (B) Quantification of HyPer ratio in A. n = 11, p = 0.001, 1 cell compared to egg, Wilcoxon matched-pairs signed rank test. (C) Schematic diagram of oocytes experiments. Immature ovarian oocytes were injected with HyPer RNA, matured with 2 *μ*M progesterone, and then pricked by a needle or laser wound activatied. (D) HyPer images of immature oocytes expressing HyPer were captured every 20 min after pricking. There is no increase in the HyPer ratio. (E) Quantification of HyPer ratio in D. n = 33, p = 0.2, 20 min compared to 0 min, paired t-test. (F) HyPer images of mature oocytes expressing HyPer were captured every 20 min after pricking. There is an increase in the HyPer ratio. (G) Quantification of HyPer ratio in F. n = 28, p < 0.0001, 20 min compared to 0 min, paired t-test. (H) SypHer images of mature oocytes expressing Sypher were captured every 20 min after pricking. (I) Quantification of SypHer ratio in H. n = 27, p < 0.0001, 20 min compared to 0 min, Wilcoxon matched-pairs signed rank test. Scale bars: (A, D, F and H) 200 *μ*m. Data are from two independent experiments. Error bars represent mean ± SEM. ***p ≤ 0.001; ****p < 0.0001; ns, not significant. See also Figure S1 and Movie S2.

To further confirm whether egg activation resulted in elevated ROS levels, we turned to an alternative ROS-detecting assay, based on Amplex Red (AR), a compound that can specifically react with H_2_O_2_ in the presence of horseradish peroxidase (HRP) to produce a fluorescent oxidated product, resorufin (Figure S1C) (Kalyanaraman et al., 2012). We injected AR into unfertilized albino eggs with or without HRP, and then we filmed the production of the fluorescent product, resorufin. We observed strong fluorescence in activated eggs injected with AR and HRP (Figure S1D and Movie S2). Taken together, these data indicate that H_2_O_2_ is generated in mature oocytes and eggs, following activation or fertilization, and that the increased ROS levels are sustained throughout early development.

### ROS production is dependent on the Ca^2+^ wave after fertilization

Previous studies have shown that sperm entry induces a Ca^2+^ wave, which activates development, cytoskeletal changes and the re-entry of the meiotic cell cycle, and subsequent initiation of the mitotic cell cycle (Nader et al., 2013). Oocyte maturation is associated with a rearrangement of Ca^2+^ signaling components, whereby oocytes acquire the ability to propagate a Ca^2+^ wave after fertilization (El-Jouni et al., 2005; Machaca, 2004). Since ROS were not induced in immature oocytes by pricking (Figures 1D and 1E), we hypothesized that Ca^2+^ signaling may act upstream of ROS production. To test this hypothesis, first we confirmed that a Ca^2+^ wave was induced by laser wounding (mimicking pricking-activation) in mature oocytes expressing R-GECO, a genetically-encoded calcium sensor (Movie S3; (Zhao et al., 2011). A similar wave was not detected in laser activated mCherry control immature or mature oocytes, nor in immature R-GECO injected oocytes (Movie S3). Notably, the Ca^2+^ wave was also induced in mature oocytes by addition of 10 *μ*M A23187, a Ca^2+^ ionophore, but not following incubation with 0.1% ethanol, the final concentration of solvent used for dissolving the ionophore (Movie S4). Importantly, addition of the Ca^2+^ ionophore also induced a 76.7% and 80% increase of HyPer ratio at 40 min and 60 min (n = 32–33, p < 0.0001, compared to control, two-way ANOVA) (Figures 2A and 2B), suggesting that activation of the Ca^2+^ wave was sufficient to induce the increase in ROS production. To determine if Ca^2+^ influx was necessary for ROS production, we incubated oocytes in the presence of the Ca^2+^ chelator EGTA (100 *μ*M) prior to activation. Extracellular Ca^2+^ has been shown to be required for fertilization in *Xenopus* (Wozniak et al., 2017). In the presence of EGTA, the Ca^2+^ wave was completely inhibited in laser-activated oocytes (Movie S5), and ROS production was also inhibited 42% and 42.2% at 40 min and 60 min after prick activation (n = 32–36, p < 0.0001, compared to control, two-way ANOVA) (Figures 2C and 2D).

**Figure 2.**
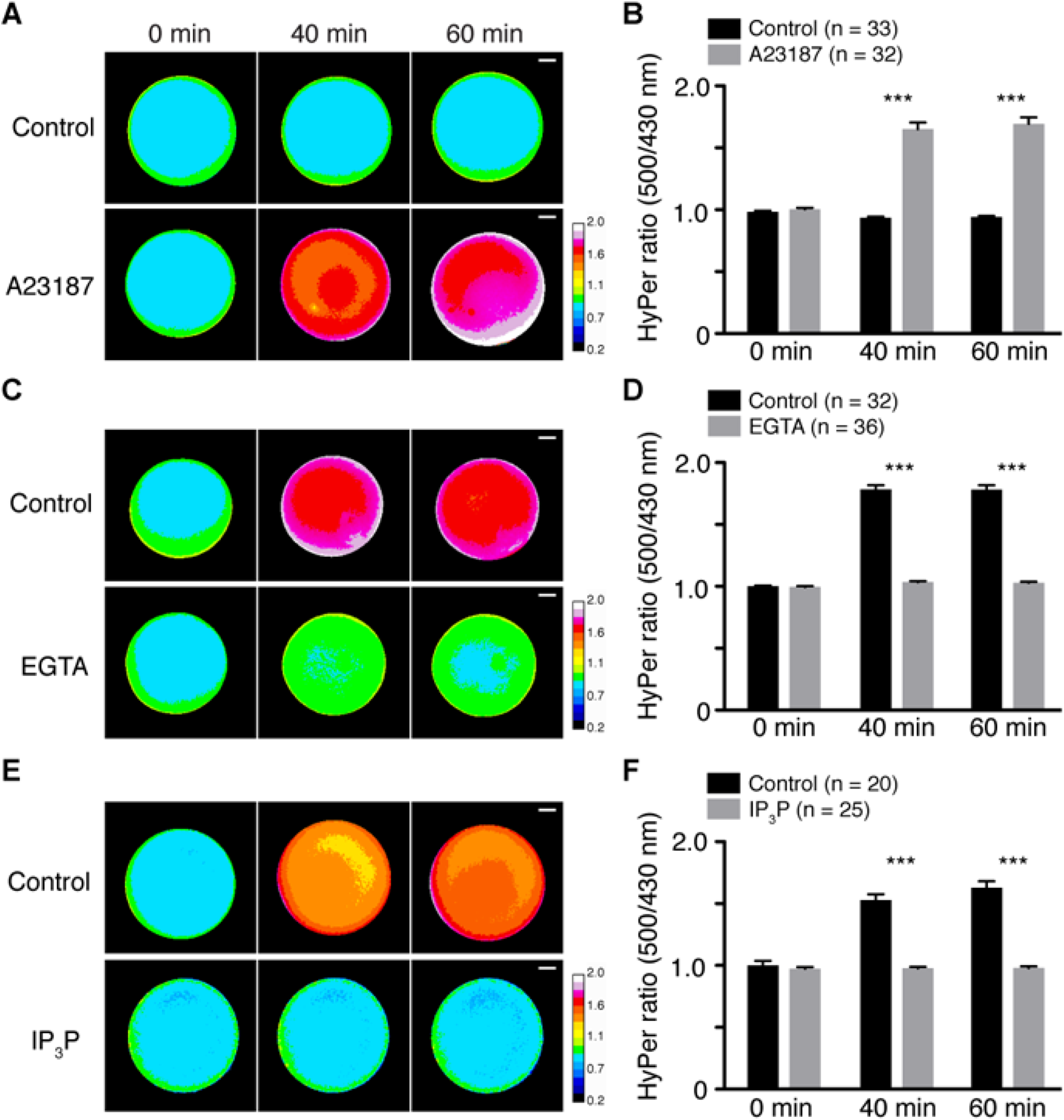
ROS production is generated downstream of Ca^2+^ signaling. (A) HyPer images of control treated or Ca^2+^ ionophore, A23187 (10 *μ*M) treated mature oocytes injected with HyPer RNA. (B) Quantification of HyPer ratio in A. n = 32–33, p < 0.0001, compared to control at 40 and 60 min. two-way ANOVA and Sidak post hoc tests. (C) Oocytes injected with HyPer RNA were matured and cultured in medium with or without EGTA (100 *μ*M), then HyPer images were taken after prick activation. (D) Quantification of HyPer ratio in C. n = 32–36, p < 0.0001, compared to control at 40 and 60 min. two-way ANOVA and Sidak post hoc tests. (E) Oocytes were injected with 20 ng of R-GECO as a control or IP3 phosphatase and HyPer RNAs, matured, and then imaged after prick activation. (F) Quantification of HyPer ratio in E. n = 20–25, p < 0.0001, compared to control at 40 and 60 min. two-way ANOVA and Sidak post hoc tests. Scale bars: (A, C and E) 200 *μ*m. Data are from three independent experiments. Error bars represent mean ± SEM. ****p < 0.001. See also Movie S3-6.

It is known that the Ca^2+^ wave in *Xenopus* zygotes after fertilization is mediated by the IP_3_ signaling pathway (Nuccitelli et al., 1993; Larabell and Nuccitelli, 1992; Runft et al., 1999). Consistently, when IP_3_ signaling was inhibited by overexpressing the IP3 phosphatase in the oocytes, the Ca^2+^ wave was completely abolished (Movie S6) and production of ROS was not detected in prick-activated matured oocytes with 36% and 39.8% reduction in ratio at 40 min and 60 min (n = 20–25, p < 0.0001, compared to control, two-way ANOVA) (Figures 2E and 2F). These results demonstrate that the fertilization/activation induced Ca^2+^ wave is both necessary and sufficient to induce ROS production in mature *Xenopus* oocytes and eggs.

### ROS production downstream of Ca^2+^ wave is mediated by mitochondria

To identify the mechanisms of ROS production downstream of Ca^2+^signaling, we used pharmacological approaches to disrupt several potential sources of ROS. Cellular ROS are mainly produced by either NOX family enzymes or the ETC in mitochondria. We found that the addition of the NOX inhibitors, 10 *μ*M diphenyleneiodonium (DPI) and 1 mM apocinin, had no effect on ROS production following prick activation in mature oocytes (n = 29–42) (Figure 3A). In addition, application of the mitochondrial complex I inhibitor, rotenone (1 *μ*M), which promotes ROS production in the forward direction and inhibits ROS production via reverse electron transport (RET) at complex I (Murphy, 2009), did not change HyPer ratios (n = 36, two-way ANOVA) (Figure 3B), suggesting that complex I is not involved in fertilization induced ROS production, neither in the forward nor reverse direction. Addition of high levels of 100 *μ*M DPI, which has also been shown to inhibit ROS production via RET from complex I (Lambert et al., 2008), also did not affect HyPer ratios in activated oocytes (data not shown), providing further evidence that the mitochondrial complex I, either in the forward or reverse direction, is not the primary source of increased ROS production following egg activation. In contrast, however, we found that incubation of the mature oocytes with 5 mM malonate, which inhibits ROS production from complex II in both the forward and reverse direction (Quinlan et al., 2012), significantly reduced ROS levels both before (52.8% decrease in ratio) and after activation (68.1–69.8% decrease in ratio) (n = 45, p < 0.0001, compared to controls at each time point, two-way ANOVA) (Figures 3C). A mitochondrial complex III inhibitor, 10 *μ*M antimycin A and complex IV inhibitor, 1 mM sodium azide, also attenuated ROS production after prick activation (23.2%−25.9% and 28.7–31.8%, respectively. n = 46 and 25, p < 0.0001, compared to control at 40 and 60 min, twoway ANOVA) (Figure 3D and 3E). We wondered whether the lack of ROS production, following addition of complex II, III and IV inhibitors might be due to depletion of ATP in the mature oocytes. Therefore, we also added the ATP synthase inhibitor, 6 *μ*M oligomycin, to the activated oocytes, but this did not affect ROS production in the oocytes (n = 42, two-way ANOVA) (Figure 3F). Furthermore, we found that addition of the ETC inhibitors, which decreased ROS production, did not decrease ATP levels in the activated oocytes (n = 3, unpaired t-test and Mann-Whitney test) (Figure S2A). In order to exclude the possibility that the ETC inhibitors affected the Ca^2+^ wave, we tested the ETC inhibitors in R-GECO expressing oocytes, and found that none of the inhibitors affected the propagation of the Ca^2+^ wave following laser wounding (n = 6, two-way ANOVA) (Figure 3G and Movie S7). Finally, we assessed whether the ETC inhibitors affected the intracellular pH of the activated oocytes expressing SypHer and we found that the SypHer ratio was not affected by activation (1 *μ*M rotenone (n = 23), 10 mM malonate (n = 19), 10 *μ*M antimycin A (n = 21), 3 mM sodium azide (n = 18) or 6 *μ*M oligomycin (n = 22) (Figure S3A-E, two-way ANOVA), suggesting that the increase in HyPer ratios were primarily due to changes in ROS levels, rather than changes in pH in the activated oocytes. In summary, these data show that mitochondria are largely responsible for ROS production following egg activation/fertilization, and interestingly, ROS production is primarily generated from complex II of the respiratory chain, and not complex I. Furthermore, the data suggest that mitochondria are not a major source of ATP production following egg activation/fertilization.

**Figure 3.**
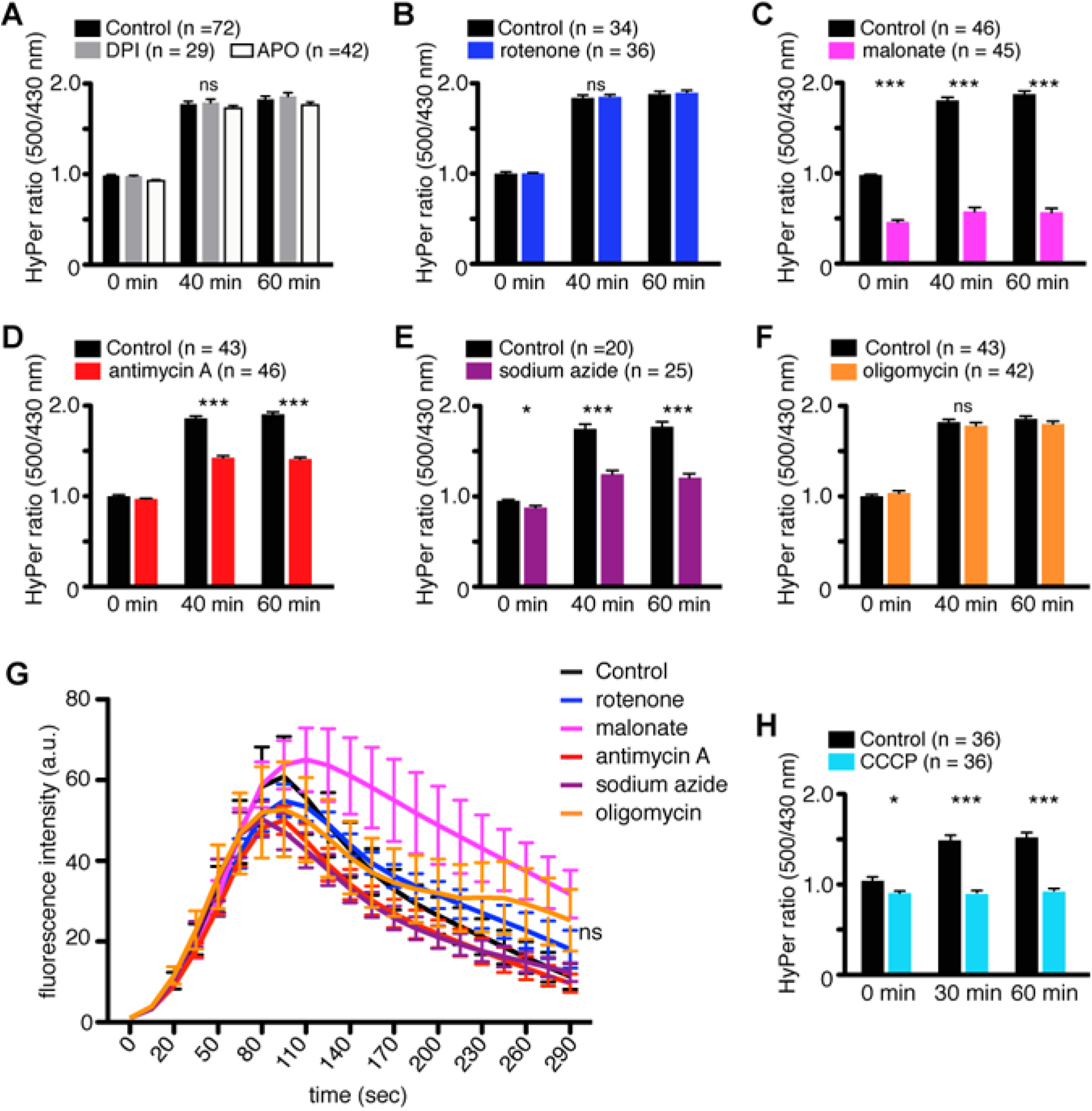
Mitochondrial inhibitors, malonate, antimycin A and sodium azide impair ROS production after activation. (A) NOX inhibitors, 10 *μ*M DPI or 1 mM apocynin had no effect on HyPer ratio in mature oocytes from HyPer transgenic females after pricking. n = 29-42, two-way ANOVA and Tukey’s post hoc tests. (B-F) HyPer ratio on oocytes treated with 1 *μ*M rotenone, n = 36 (B), 5 mM malonate, n = 45, p < 0.0001, compared to control at each time point, two-way ANOVA and Sidak post hoc tests (C), 10 *μ*M antimycin A, n = 46, p < 0.0001, compared to control at 40 and 60 min, two-way ANOVA and Sidak post hoc tests (D), 1 mM sodium azide, n = 25, p < 0.0001, compared to control at 40 and 60 min, two-way ANOVA and Sidak post hoc tests (E) and 6 *μ*M oligomycin, n = 42 (F). (G) Ca^2+^ wave was measured by fluorescent intensity of R-GECO in laser-activated mature oocytes. Ca^2+^ wave was not affected by any of the mitochondrial inhibitors. Each treatment, n = 6, two independent experiments. two-way ANOVA and Tukey’s post hoc tests. See also Movie S7. (H) HyPer ratio on oocytes treated with 2 *μ*M FCCP. n = 36, p < 0.0001, compared to control at 40 and 60 min, two-way ANOVA and Sidak post hoc tests. Data for A-F and H are from three or four independent experiments. Error bars represent mean ± SEM. ns, not significant; ****p < 0.0001.

To confirm whether the mitochondria are indeed responsible for ROS production after egg activation/fertilization, we employed a mitochondrial targeted HyPer construct (mitoHyPer) to more specifically detect ROS produced in mitochondria (Malinouski et al., 2011). Immature oocytes were injected with mitoHyPer RNA, and the oocytes were then matured and activated by pricking in the presence of the ETC inhibitors. Although the increase in mitoHyPer ratios were less apparent than that seen with the cytosolic version of HyPer, similar effects of inhibitors on ROS production were observed using mitoHyper, in that 5 mM malonate (n = 46, 19.8%, 34.6% and 35.7% at 0, 40 and 60 min), 10 *μ*M antimycin A (n = 42, 14.5% and 16.5% at 40 and 60 min) and 1 mM sodium azide (n = 48, 13.1% and 14.9% at 40 and 60 min) decreased ROS production significantly in activated oocytes (p < 0.0001, compared to control, two-way ANOVA), while rotenone (n = 46) and oligomycin (n = 40) did not (Figure S3F-J). These findings suggest that, while ROS production is primarily generated in the mitochondria, the bulk of the ROS generated within the mitochondria rapidly diffuses into the cytoplasm.

ROS production in mitochondria is known to be regulated by the amplitude of the mitochondrial membrane potential (ΔΨm) (Andreyev et al., 2005). To examine if ROS production in oocytes depends on the ΔΨm, oocytes expressing HyPer were treated with the mitochondrial uncoupler, carbonyl cyanide 3-chlorophenylhydrazone (CCCP). As shown in Figure 3H, ROS production was significantly reduced by CCCP treatment when compared with controls (39.5% and 39.1% decrease at 30 and 60 min, n = 36, p < 0.0001, compared to control at 30 and 60 min, two-way ANOVA). Taken together, these data suggests that mitochondria are responsible for the generation of ROS downstream of Ca^2+^ after fertilization/activation.

### Ca^2+^ wave directly mediates ROS production by mitochondria via MCU

Previous reports have shown that mitochondria uptake calcium from the cytosol via the mitochondrial calcium uniporter (MCU) (Duan et al., 2007; Rharass et al., 2014; Szabadkai and Duchen, 2008; Xu and Chisholm, 2014). We therefore wondered whether MCU-mediated calcium import was essential for the ROS production following fertilization. To test this hypothesis, we injected oocytes with 400 pM ruthenium red (RuR), an inhibitor of MCU, along with HyPer mRNA. Indeed, oocytes injected with RuR and HyPer mRNA failed to produce ROS after pricking (29.1–30.5% decrease, n = 23–24, p < 0.0001, compared to control at each time point, Mann-Whitney test) (Figure 4A and 4B). In order to confirm that the inhibition of ROS production in oocytes injected with RuR was specific to MCU, we explored an independent way of inhibiting MCU. In this regard, it had been previously shown that the mitochondrial calcium uniporter functions as a tetramer. Furthermore, Raffaello and colleagues (Raffaello et al., 2013) showed that *mcub* (originally named as *CCDC109B*), encodes an MCU related protein that acts as an endogenous dominant negative of MCU activity, disrupting the ability for MCU to form calcium-permeable functional tetrameric channel. To exploit this inhibitory property of *mcub*, we injected mRNA encoding *X. tropicalis mcub* RNA with HyPer RNA and found that overexpressing *mcub* reduced ROS production in the prick activated mature oocytes (9.4% and 11.5% decrease, n = 24, p < 0.01, compared to control at 30 min and p < 0.001 at 60 min, two-way ANOVA) (Figure 4C and 4D). Notably, the Ca^2+^ wave after laser activation was not affected by overexpression of *mcub* (Movie S8). These data show that the increase in Ca^2+^ after fertilization enters the mitochondria via the mitochondrial calcium uniporter, and this results in an increase in mitochondrial ROS production.

**Figure 4.**
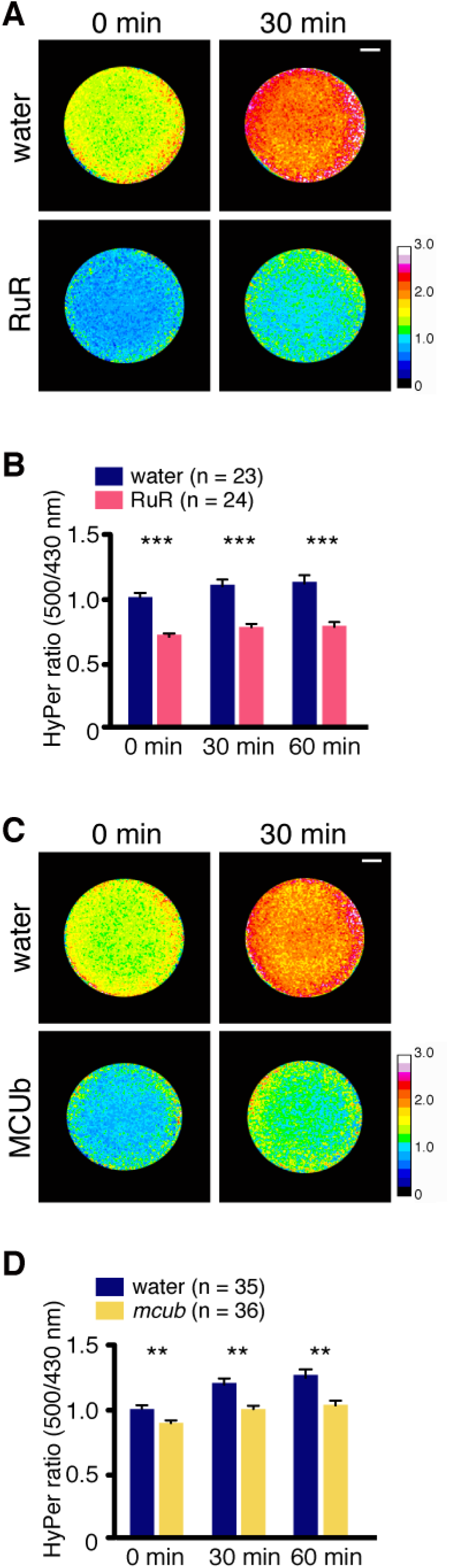
Ca^2+^ mediates ROS production by mitochondria via MCU. (A) HyPer images of oocytes injected with 40 nl of containing 0.4 pM RuR (0.1 *μ*M final conc.), MCU inhibitor and 20 ng HyPer RNA. (B) Quantification of HyPer ratio in A. ROS production was reduced by MCU inhibition. Two independent experiments. ****p < 0.0001, compared to control at each time point. Mann-Whitney test. Error bars represent mean ± SEM. (C) HyPer images of oocytes injected with 20 ng of RNA for dominant negative MCU, MCUb and 20 ng HyPer RNA compared to 20 ng HyPer RNA injected control. (D) Quantification of HyPer ratio in C. Inhibition of MCU by overexpression of MCUb impairs ROS production. Two independent experiments. **p < 0.01, ***p < 0.001, compared to control at t30 and t60, two-way ANOVA and Sidak post hoc tests. See also Movie S8. Scale bars: (A and D) 200 *μ*m.

### Mitochondrial ROS inhibition causes cell division arrest during the early cleavage stages of the embryo

To elucidate the role of ROS produced by mitochondria in early development, onecell stage embryos were treated with various mitochondrial inhibitors toward the end of the first cell cycle. Figure 5A shows phenotypes of treated embryos observed at the 32-cell stage. Embryos treated with 1 *μ*M rotenone stopped dividing at the 2-cell stage (100%, n = 92). 10 mM malonate and 3 mM sodium azide also caused cell division arrest at the 4 or 8-cell stage (100%, n = 40–58). Embryos treated with 10 *μ*M antimycin A developed normally until the 64-cell stage (97.6%, n *=* 83), however, their division rates were delayed and ultimately underwent embryonic death at the blastula stage (100%, n = 83). Embryos treated with 6 *μ*M oligomycin developed normally up to the blastula stage (n = 88) (Figure 5A).

**Figure 5.**
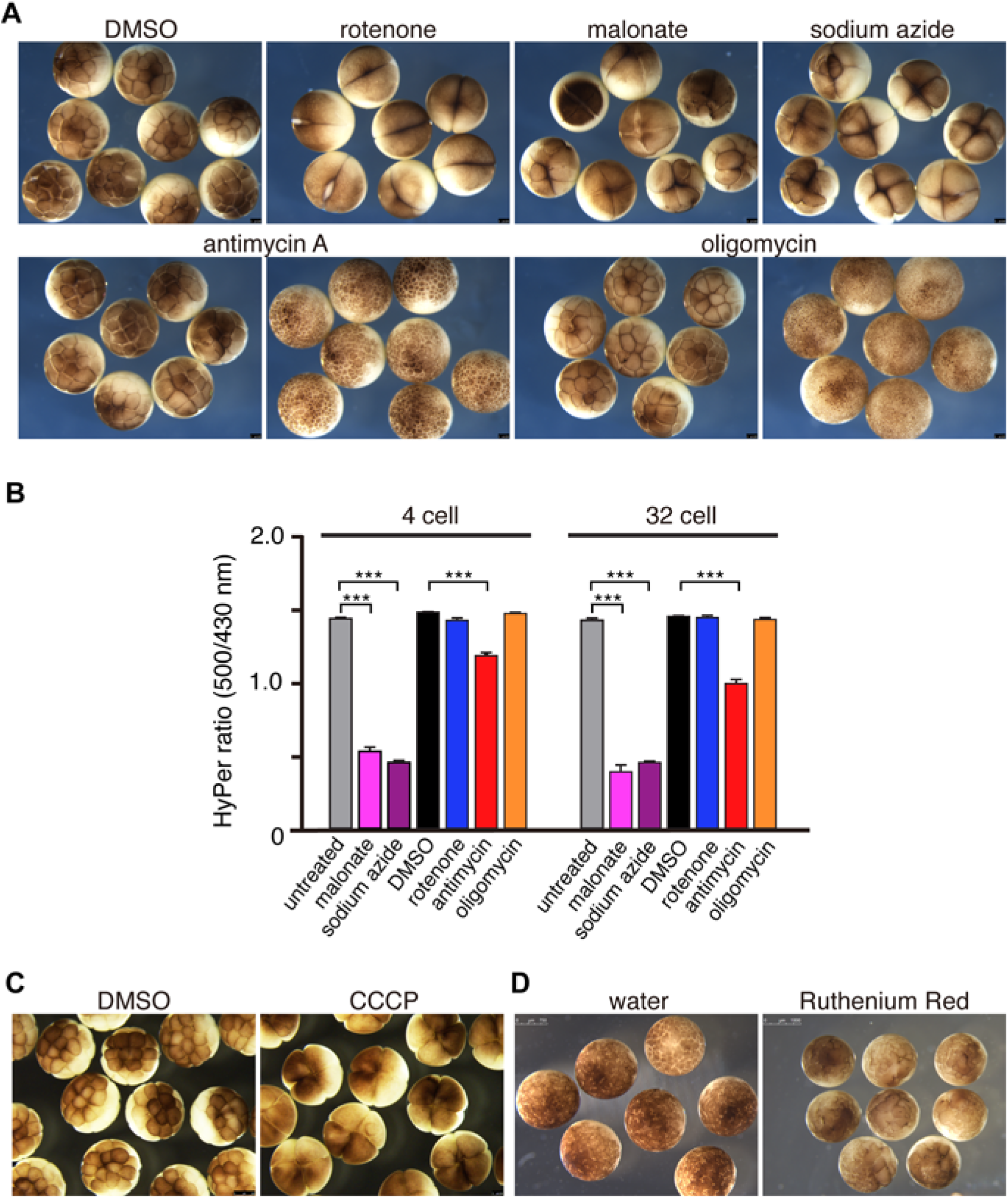
Inhibition of mtROS induces cell division arrest in *Xenopus* early development. (A) Embryos were treated at the one-cell stage with 0.1% DMSO (n = 36), 1 *μ*M rotenone (n = 92), 10 mM malonate (n = 40), 3 mM sodium azide (n = 58), 10 *μ*M antimycin A (n = 83) and 6 *μ*M oligomycin (n = 88). Embryos treated with rotenone, malonate and sodium azide were arrested at the 4 to 8 cell stage. By the blastula stage embryos treated with antimycin A were arrested. (B) 1-cell stage embryos obtained from females expressing HyPer were treated with mitochondrial inhibitors, and HyPer ratio was measured at the 4-cell and 32-cell stages. n = 12-13; ****p < 0.0001. two-way ANOVA and Tukey’s multiple comparisons tests. Error bars represent mean ± SEM. (C) Embryos were treated with 0.1% DMSO or 2 *μ*M CCCP (n = 66). (D) Embryos were injected with water as a control and 50 pM RuR into one cell at the 2-cell stage. Injection of RuR induced cell cycle arrest (n = 56). Pictures in A, C and D are representative of at least three independent experiments.

To confirm whether these phenotypes correlated with reduction of ROS production, 1-cell stage embryos obtained from transgenic females expressing HyPer were treated with the various mitochondrial inhibitors to monitor ROS levels. We observed an increase in HyPer ratio in untreated (n = 12) and DMSO (n = 13) control embryos at the 4-cell and 32 cell stages (Figure 5B). As seen in oocytes, malonate (n = 11) and sodium azide (n = 12) significantly reduced ROS level 62.9% and 67.7%, respectively, in 4-cell stage embryos in which cell division was arrested (p < 0.0001, compared to untreated control, two-way ANOVA). Antimycin A, which caused cell-cycle delay and eventual arrest only at the blastula stage, only slightly reduced ROS production (20.3% at 4-cell, n = 12, p < 0.0001, compared to DMSO control, two-way ANOVA). Oligomycin, an ATP synthase inhibitor, had no effect on either the cell division rates nor ROS levels (n = 11). Although rotenone induced rapid cell division arrest, ROS reduction in these embryos was not observed (n = 12). Given that rotenone has been shown to affect actin dynamics through modification of Rho-GTPase (M. Sanchez et al., 2008), which are known to be necessary for cytokinesis (Drechsel et al., 1997), we interpret this early cell cycle division defect as an inhibition of cytokinesis, rather than inhibiting of the cell cycle oscillator. ATP levels in the embryos treated with the various inhibitors were also measured and we confirmed that there were no significant reductions in ATP levels in all inhibitor-treated embryos (n = 6, unpaired t-test) (Figure S2B), and thus the cell cycle arrest cannot be due to a decrease overall ATP levels in the early embryos. Consistent with ROS reduction in oocytes treated with CCCP (Figure 3H), embryos treated with 2 *μ*M CCCP also exhibited cell division arrest at the 4-cell stage (100%, n = 66) (Figure 5C). Injection of 50 pM RuR, the MCU inhibitor, that caused reduction of ROS in oocytes (Figure 4A) also induced cell division defects at the injection site (100%, n = 56, Figure 5D). Taken together, these data suggest a strong link between ROS production from mitochondria and progression of the cell cycle arrest.

We then asked whether the defect of cell division could be rescued by addition of either H_2_O_2_ or menadione, but neither were able to rescue the cell division arrest caused by malonate or sodium azide (data not shown). This is likely due to the difficulty in reconstituting the correct level of ROS and its dynamics, via simple addition of these oxidants. However, given that malonate and sodium azide are reversible inhibitors of mitochondrial complex II and complex IV, respectively, we tested whether cell division and ROS production could be restored after the removal of these inhibitors. Thus, we treated fertilized embryos with the inhibitors, and once they had arrested at the 4-cell stage, we transferred the embryos into fresh medium without the inhibitors. The removal of inhibitor led to the re-initiation of the cell cycle in 48% of embryos treated with malonate and 93% of embryos treated with sodium azide (n = 42-44) (Figure 6A). Remarkably, most of the rescued embryos then developed to the swimming tadpole stage. Using embryos expressing HyPer, we confirmed that ROS production was restored within 30 minutes after the removal of inhibitors (Figure 6B and Movie S9). After treatment of inhibitors (0 min), HyPer ratios were 49.1% in malonate treated embryos and 33.8% in sodium azide treated embryos compared to control embryos, but they recovered to 84.9% and 78.9% within 40 min after removal of inhibitors, respectively (n = 36). Taken together, these data confirm that the cell division defects observed in the embryos treated with malonate and sodium azide were primarily caused by the reduction in ROS levels and that the cell cycle is able to re-initiate once ROS levels are allowed to be restored following inhibitor removal.

**Figure 6.**
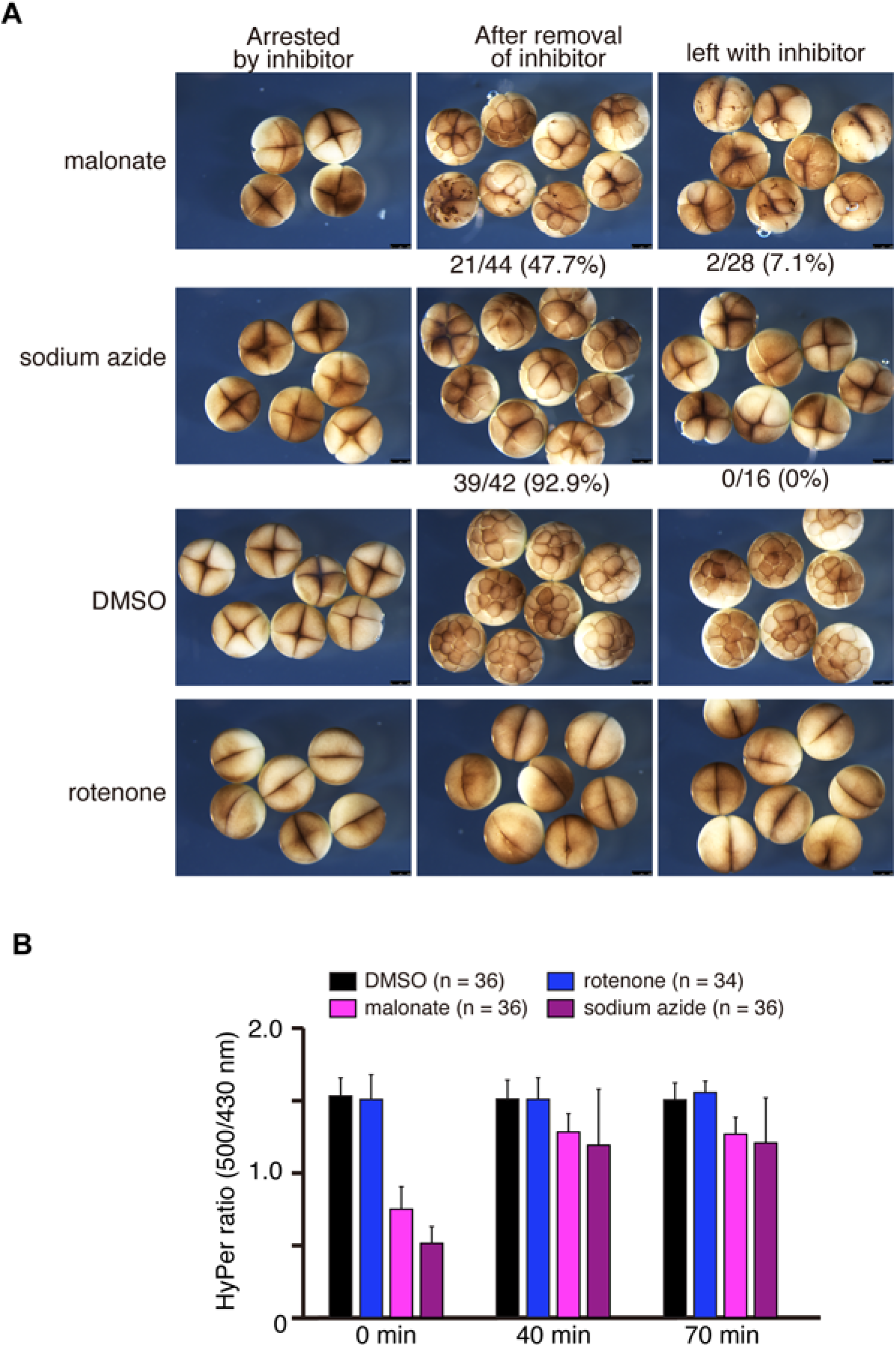
Removal of inhibitors restores cell division and ROS production. (A) Embryos were treated with inhibitors, 10 mM malonate, 3 mM sodium azide and 1 *μ*M rotenone for 40 min until cells stopped dividing (left column), then transferred to a dish containing fresh medium without inhibitors. Embryos transferred after treatment with malonate and azide retrieved cell division and divided to 8–16 cell stages (middle), but not in medium with inhibitors (right). Pictures are representative of three independent experiments. (B) Fertilized transgenic embryos expressing HyPer were treated with inhibitors for 40 min, and imaged shown as 0 min. Then embryos were transferred to fresh medium without inhibitors, and imaged at 40 and 70 min. n = 34–36, three independent experiments. Error bars represent mean ± SD. See also Movie S9.

### Cdc25C-dependent cell cycle progression is regulated by ROS

Next we sought to better understand the molecular mechanisms by which ROS may regulate the cell cycle during the early cleavage stages of *Xenopus* embryos. Given that H_2_O_2_ is a potent in inhibitor of protein tyrosine phosphatases through the reversible oxidation of their catalytic cysteine residues (Lee et al., 2002; Salmeen et al., 2003), we decided to focused on the Cdc25C phosphatase, a key regulator of the cell cycle that, in yeast, had been shown to be redox sensitive (Rudolph, 2005; Seth and Rudolph, 2006). We therefore endeavored to ask whether Cdc25C might be a target of ROS during the early *Xenopus* mitotic cycles, especially given that *Xenopus* Cdc25C is present at constant levels during early *Xenopus* development (Kumagai and Dunphy, 1992).

Cdc25C activates the cyclin B-Cdk1 complex to induce entry into mitosis by dephosphorylating the inhibitory-phosphorylated Thr 14 and Tyr 15 residues in Cdk-1 (Perdiguero and Nebreda, 2004). Cdc25C itself is also regulated by phosphorylation, and it has been shown that there are at least five sites in Cdc25C, whose phosphorylation correlates with increased phosphatase activity. In contrast, hypophosphorylation of those five sites and phosphorylation on Ser 287 (Ser216 in human) is associated with inactive Cdc25C (Kumagai et al., 1998; Matsuoka et al., 1998; Peng et al., 1997; Y. Sanchez et al., 1997; Zeng et al., 1998). To examine Cdc25C phosphorylation, we performed immunoblots using *anti-Xenopus* Cdc25C antibodies that recognize both the active and inactive forms of Cdc25C. Embryos treated with the various mitochondrial inhibitors were collected every 15 minutes and subjected to Western blot analysis. Notably, extracts from DMSO control treated embryos exhibited oscillating hyper and hypophosphorylation states of Cdc25C, depending on the stage of the cell cycle, correlating with cycling Cdc25C activity (Figure 7A). Inhibitors that had no effect on ROS production, such as 1 *μ*M rotenone and 6 *μ*M oligomycin, did not change the oscillating state of Cdc25C phosphorylation during the cell cycle, suggesting that these inhibitors do not affect the cell cycle oscillator. In contrast treatment with the inhibitors 10 mM malonate and 3 mM sodium azide, which cause a reduction in ROS levels, resulted in either a delay or total inhibition in the cycling Cdc25C hyper - hypophosphorylation oscillations in early embryos (asterisks in Figure 7A), suggesting that reduction in ROS levels affected the cycling activation / deactivation pattern of Cdc25 during the cell cycle.

**Figure 7.**
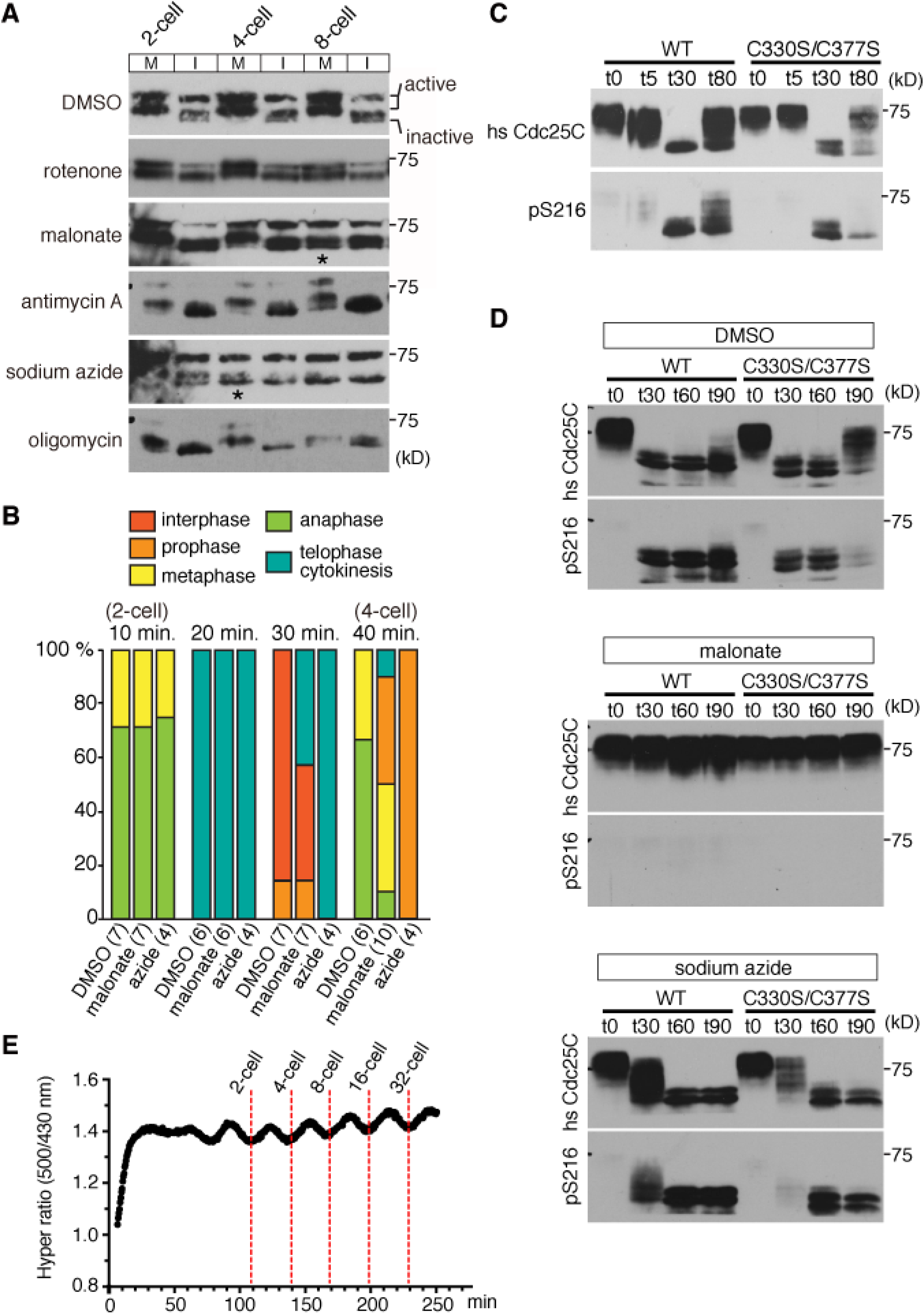
Inhibition of mtROS causes misregulation of Cdc25C activity resulting in mitotic arrest. (A) Immunoblots of *Xenopus* Cdc25C in embryos treated with mitochondrial inhibitors. 10 mM malonate and 3 mM sodium azide, which reduced ROS generation, caused misregulation of Cdc25C (* in the blots). (B) Embryos were treated with inhibitors at the 1-cell stage (90 min after fertilization) and fixed every 10 min for immunohistochemistry. The staining for α-Tubulin, Lamin B1 and DNA were used to identify the phase of cell-cycle. The numbers in parentheses indicate numbers of embryos examined. A defect in mitotic entry was observed in embryos treated with malonate and sodium azide. Data are from two to three independent experiments. (C) Immunoblots of *Xenopus* Cyclin B2 in embryos treated with 10 mM malonate or 3 mM sodium azide. Cyclin B2 degraded before cell cycle arrest (* in the blots), but its accumulation seemed to be impaired afterward. (D) Immunoblots of *Xenopus* Cdc25C, Cdk-Y15, pan-Cdk, Cyclin B2 and α-Tubulin in activated eggs treated with 0.1% DMSO, 10 mM malonate or 3 mM sodium azide at 0, 5, 30 and 80 min after prick activation. (E) Immunoblots detecting human Cdc25C and human Cdc25C-pS216 in activated mature oocytes. Oocytes injected with human Cdc25C WT or C330S/C377S RNA were treated with DMSO or mtROS inhibitors, prick-activated and extracted at 0, 30, 60 and 90 min. All blots are representative of at least three independent experiments. (F) Plots of measurement of HyPer ratio shown in black (left Y axis) and the raw fluorescence intensities at 500 nm of HyPer and YFP embryos shown in the plot as green and yellow lines, respectively (right Y axis). Embryos were imaged in every 30 sec from at the beginning of fertilization (0 min) throughout the cleavage stage in Movie S10. Data are representative from at least two independent experiments. See Movie S11. (G) YFP(500 nm)/CFP (430 nm) ratio of embryos expressing HyPer (green) or SypHer (black and pink) obtained by nuclear transplantation of in vitro matured oocytes. Embryos were imaged in every 30 sec when it started dividing (0 min) throughout the cleavage stage. Data are representative from at least two independent experiments (n = 2–4).

Rotenone inhibited cytokinesis but had no effect on neither ROS production nor cell cycle oscillator, as evidenced by the continued cycling of Cdc25C hyper-hypo phosphorylation pattern (Figures 5A, 5B and 7A). It is known that inhibition of cytokinesis does not affect the nuclear cell cycle oscillator in early *Xenopus* embryos (Gerhart and Kirschner, 1984). Thus cytokinesis cannot be used as sole measure of whether the cell cycle oscillator is operational or not in early frog embryos, as evidenced by the lack of cytokinesis in embryos treated with rotenone, yet these embryos still show oscillating Cdc25 hyper-hypo phosphorylation pattern.

To investigate which phase of the cell cycle was affected by the various treatments, immunohistochemistry was carried out on embryos using antibodies against α-Tubulin and Lamin B1, alongside sytox DNA staining to visualize the chromosomes. As shown in Figure 7B, control embryos proceeded to anaphase of the second cell cycle at 40 min after treatment. On the other hand, embryos showing cell cycle arrest at 40 min after treatment with malonate or sodium azide had nuclei and microtubules typically seen in prophase. Embryos treated with rotenone proceeded to anaphase similarly to control embryos (data not shown), confirming that the phenotype caused by rotenone is not due to a defect in the nuclear cell cycle. Thus embryos treated with the mtROS inhibitors (malonate and sodium azide) lost Cdc25C cycling activity and failed to proceed through mitosis resulting in cell cycle arrest.

It has been known that degradation of Cyclin B is critical for mitotic exit. Therefore, we wished to ask whether Cyclin B degradation was affected in embryos treated with the mtROS inhibitors. As shown in Figure 7C, Cyclin B degraded at interphase in control embryos. In embryos treated with the mtROS inhibitors, Cyclin B degradation was observed as expected in interphase before cell cycle arrest at 4-cell in azide and at 8-cell in malonate (asterisks in Figure 7C). However, after cell cycle arrest the accumulation of Cyclin B required for mitotic re-entry was not detected. This observation indicates that failure of mitotic arrest caused by mtROS inhibitors is not due to disregulation of Cyclin B degradation, although we do find a disregulation in Cyclin B re-accumulation in the subsequent cell cycle. Together, these findings suggest that mtROS has a role in regulating the oscillation of Cdc25C activity required for cell cycle progression.

To determine whether mtROS have an impact on Cdc25C phosphatase activity, the phosphorylation state of Tyr 15 of Cdk1, a direct target of Cdc25C, was examined by immunoblot using an anti-Cdk1-Y15 antibody (Tsai et al., 2014). To examine how Cdc25C activity is regulated by mtROS, unfertilized eggs were activated by pricking and collected at different time point. As shown in Figure 7D, endogenous Cdc25C was hyperphosphorylated before activation (t0) and was enzymatically active since phosphorylation of Cdk1 at Tyr 15 was not detected. 30 min after prick-activation (t30) control mature oocytes exhibited hypophosphorylation, and coincident with this, its specific substrate (Cdk1-Tyr15) was phosphorylated, suggesting diminished Cdc25C activity (Figure 7D). However, under condition where mtROS were inhibited by either malonate or sodium azide, Cdc25C remained enzymatically active after 30 minutes as suggested by undetectable Cdk1-Tyr 15 phosphorylation (Figure 7D). Intriguingly, sodium azide, which is the more potent inhibitor of the cell cycle, displayed continued Cdc25 activity and/or Wee1 inactivity, even after 80 minutes, when control oocytes showed recovery of Cdk1-Tyr15 phosphorylation. In addition, activated oocytes treated with malonate, which delays progression through the cell cycle prior to complete inhibition, exhibited an intermediate level Cdk-Tyr15 phosphorylation, suggesting continued Cdc25 activity after 80 minutes and/or diminished Wee1 activity, relative to the control oocytes (Figure 7D). Consistent with our findings in embryos (Figure 7C), Cyclin B degradation still occurred normally by t30 in the activated oocytes treated with malonate or sodium azide, but both inhibitors either delayed (malonate) or inhibited (sodium azide) the recovery of cyclin B levels at t80 (Figure 7D). These data indicate that mtROS are involved in the rapid down-regulation of Cdc25C activity at the meiotic exit and, in its absence, Cdc25 activity remains high, especially under continual sodium azide treatment.

Cdc25 has been proposed to be regulated by oxidation of the conserved cysteines in the catalytic domain (Rudolph, 2005; Seth and Rudolph, 2006). To test whether these cysteine residues are involved in the dynamic post-translational regulation of Cdc25C, we utilized the oocytes system to overexpress human Cdc25C containing mutations in the two critical redox sensitive cysteine residues (C330S and C377S) (Savitsky and Finkel, 2002). Monitoring the human Cdc25C protein also allowed us to assess the dynamics of the phosphorylation state of serine 216, which is inhibitory to its subcellular localisation and activity (Takizawa and Morgan, 2000), as the available anti-Cdc25C (phospho S216) antibody reacts with the human isoform, but not the endogenous *Xenopus* Cdc25 protein at its homologous position at serine 287. Activation of Cdc25C induced by progesterone has been shown to be important for oocyte maturation and overexpression of Cdc25C induces precocious oocyte maturation in *Xenopus* (Perdiguero and Nebreda, 2004; Gautier et al., 1991). Indeed we noticed that injection of 5 ng wild type human Cdc25C RNA induced oocyte maturation without progesterone treatment. Since overexpression of mutant human Cdc25C RNA did not inhibit maturation induced by progesterone and coinjection of human Cdc25C mutant RNA (1 ng to 10 ng) with 5 ng wild type human Cdc25C RNA did not prevent the precocious maturation by the wild type Cdc25C, it is unlikely that mutant human Cdc25C acts as a dominant negative. We also confirmed that injection of 500 pg mutant RNA had no effect on embryonic development when it was injected at the 1-cell stage (data not shown). For our experiments, we injected 100 pg of wild type or mutant human Cdc25C RNAs, which did not induce precocious oocyte maturation nor did they disrupt maturation induced by progesterone. We found that both the wild type (WT) and the C330S/C377S mutant form of human Cdc25 were hyper-phoshorylated in the oocytes prior to prick activation. Furthermore, at t0, no detectable phosphorylation at S216 was observed, suggesting that both the WT and mutant form had the post-translational hallmarks of active Cdc25 (t0 in Figure 7E, top DMSO control treated oocytes). 30 min after prick activation, both the WT and mutant Cdc25 became fully hypophosphorylated and both forms contained the inactive S216 phosphorylation state (t30 in Figure 7E, top DMSO control treated oocytes). Inhibiting mtROS production, following oocyte activation, by treating them with malonate resulted in the retention of the hyper-phosphorylated WT and mutant forms of Cdc25, as well as the sustained lack of phosphorylation of the inhibitory S216 site, suggesting that inhibition of mtROS via malonate, resulted in the retention of post-translational modification of Cdc25 associated with a sustained active state (Figure 7E, middle malonate treated oocytes). Furthermore, inhibiting mtROS with sodium azide resulted in a delay in the change from hyper to hyperphosphorylation state of the WT and mutant forms of Cdc25, such that at t30, there still remained significant amounts of hyperphosphorylated Cdc25, while DMSO treated oocytes at t30 had no hyperphosphorylated Cdc25 left, and interestingly, the mutant form was more delayed in acquiring its inhibitory phosphorylation at S216 than the WT form at t30, suggesting that the rate of de-activation in the mutant was slower when Cdc25 lacked the two ROS sensitive cysteines (Figure 7E, bottom sodium azide treated oocytes). These data suggest that mtROS facilitate the inactivation of Cdc25C through its ROS-sensitive cysteine residues.

### ROS oscillate with the cell cycle

Having seen that inhibiting ROS production leads to cell cycle arrest, as well as the loss of the oscillatory phosphorylation and activity states of Cdc25, we hypothesized that ROS levels might also oscillate during the early *Xenopus* embryo, coincident with the cell cycle. Thus we returned to HyPer imaging of early embryos and began imaging the embryos in greater detail. We found that ROS levels do fluctuate during the early cell cycles, and the peaks of ROS levels coincide with cytokinesis of each cycle, and the troughs with interphase between divisions (Figure 7F, black line, Movies S10 and S11). We also imaged embryos expressing YFP maternally, and fluorescence intensity at 500 nm was measured every 30 sec. As shown in Figure 7F, while HyPer signal (green line) showed oscillatory fluctuation, YFP (yellow line) did not. To eliminate the possibility that an oscillating change in pH may be responsible for the oscillating HyPer pattern, we generated embryos expressing SypHer. While we observed a clear oscillation pattern in HyPer ratios (green line), we failed to observe oscillating SypHer ratios in the embryos (pink and black lines) (Figure 7G). These results strongly suggest that ROS levels oscillate through the cell cycle, thus explaining why the cell cycle arrest could not be simply rescued by addition of H_2_O_2_ or menadione, as these treatments would not have reconstituted the oscillating patterns of ROS, necessary for restoring the cell cycle.

## DISCUSSION

We have previously shown that tail amputation in *Xenopus* tadpoles induces a sustained increase in ROS levels throughout tail regeneration, and these increased levels are required for tail regeneration (Love et al., 2013). Intriguingly, we show here that fertilization also triggers increased ROS levels, which are sustained, yet oscillating with the cell cycle, during early embryonic development. Given that the events triggered by fertilization can be mimicked by prick activation or laser wounding, one might consider fertilization as a sort of injury, which triggers sustained ROS production and cell cycle progression and development, in the same way that tail amputation triggers sustained ROS production, which is essential for cell proliferation, growth factor signaling and, ultimately, appendage regeneration. Indeed, we also know that both tissue injury and fertilization induce a rapid Ca^2+^ wave (Soto et al., 2013), which in the context of fertilization, is necessary and sufficient for ROS production, via the Ca^2+^ uniporter, MCU. Intriguingly, however, we find that the source of ROS production is different in both scenarios. While NADPH oxidase activity is the primarily source of ROS production following tail amputation (Love et al., 2013), ROS production following fertilization or egg activation is primarily dependent on the mitochondrial ETC, more specifically complex II, III and IV. Although we have used DPI as a potent NADPH oxidase inhibitor, DPI has also been shown to inhibit the FMN coenzyme of complex I (Majander et al., 1994) and ROS production from complex I via RET (Lambert et al., 2008). In our studies, neither high concentration of DPI (100 *μ*M) nor rotenone affected ROS production in oocytes or embryos, indicating that complex I is not responsible for ROS production in early embryos. Thus we find that complex I, the site traditionally viewed as the major source of mitochondrial ROS production (Mailloux, 2015; Murphy, 2009), is not involved in ROS production in the early frog embryos, suggesting the mitochondria in early embryos do not behave in a canonical manner. This is further supported by our finding that inhibiting the mitochondrial ETC does not decrease ATP levels in the early embryos. It is thus fascinating to speculate whether mitochondria in early embryos are primarily used as signaling organelles, via ROS production, rather than ATP producing organelles. The use of mitochondria-targeted antioxidants may provide further information for understanding the mechanism of ROS production by mitochondria during early embryogenesis.

Our previous study on the role of ROS production during appendage regeneration showed that inhibiting ROS production resulted in a significant decrease in proliferation following tail amputation (Love et al., 2013). However, in that study, we also found that both Wnt and FGF signaling were attenuated when ROS production was inhibited. Given that cell proliferation is often controlled via growth factor signaling, it was not clear from that study whether the cell proliferation defect was a consequence of attenuated growth signaling or whether ROS levels affected growth factor signaling and cell proliferation independently. This study provides evidence that ROS is capable of impacting cell proliferation independently from their effects on growth factors signaling. This is because the early cell cycle progression in frog embryos is not dependent growth factor signaling. Indeed growth factor signaling does not play a role in development until the mid-blastula stage, when zygotic transcription is initiated (Newport and Kirschner, 1982; Zhang et al., 2013), yet we find that attenuating ROS levels affects the cell cycle at the cleavage stages. Intriguingly, after a burst of ROS production following fertilization, we find that ROS levels then oscillate with the cell cycle, and that the elevated, oscillating ROS levels are necessary for the cell cycle to progress. We were unable to rescue cell cycle arrest caused by mtROS inhibitors by adding H_2_O_2_ or menadione. This is likely due to the fact that ROS levels oscillate during the cell cycle, an aspect which was not reproduced adequately in the attempted rescue experiments.

The control of the cell cycle in *Xenopus* early embryos involves positive and negative feedback loops, which constitute a bistable mitotic trigger (Thron, 1996). For generation of robust bistability, ultrasensitivity of Cdc25C has been suggested (Trunnell et al., 2011). In this study, we have shown that the conserved cysteine residues in Cdc25C help mediate a quick inactivation of Cdc25C, suggesting that ROS may supply Cdc25C with the necessary ultrasensitivity for robustness suggested by Trunnell and colleagues. Although here we have focused on the redox-sensitive Cdc25C, future studies will strive to identify other redox-sensitive proteins and how they utilize the fertilization-induced, oscillatory ROS production to ensure successful progression through early embryonic development.

## MATERIALS AND METHODS

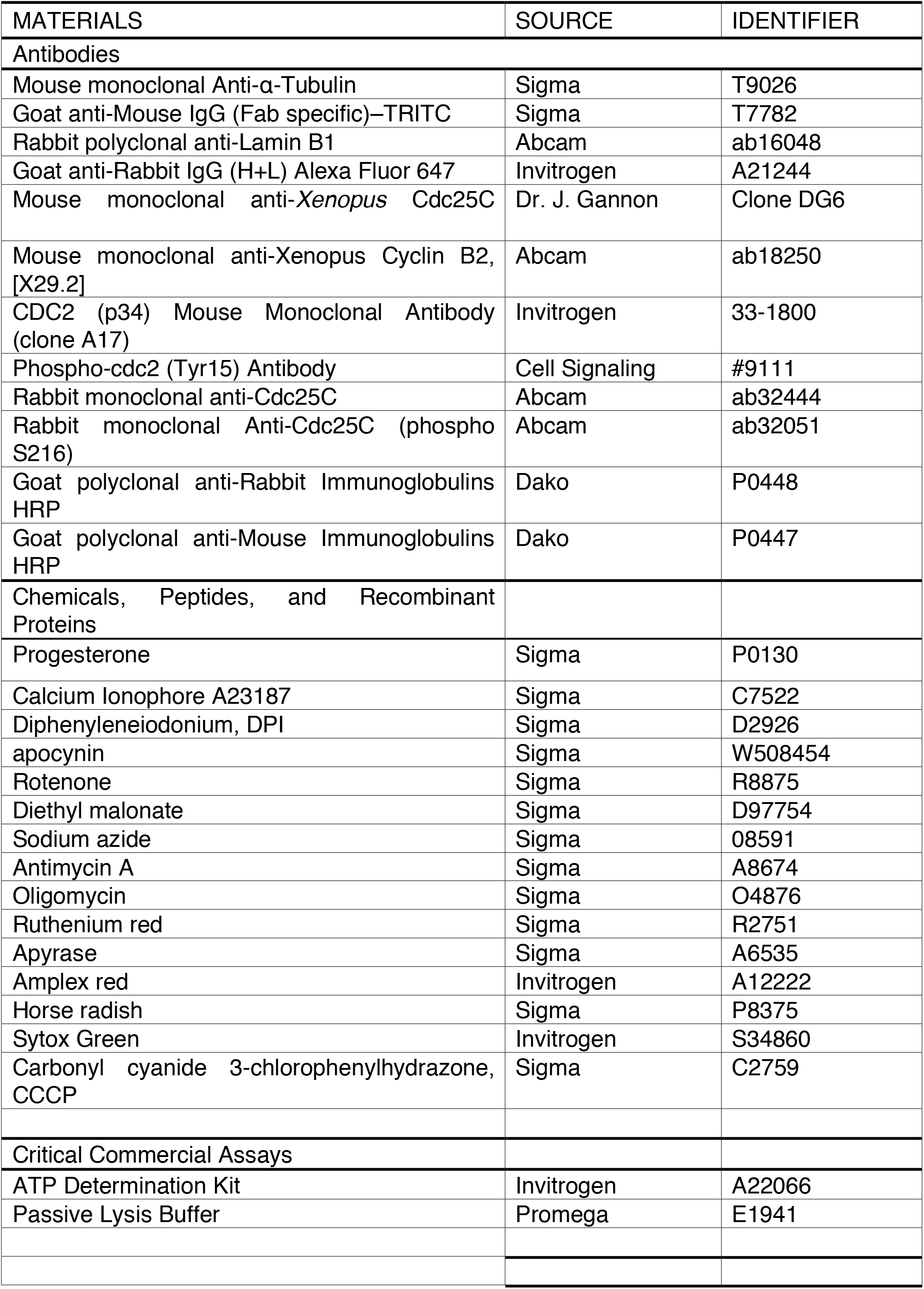
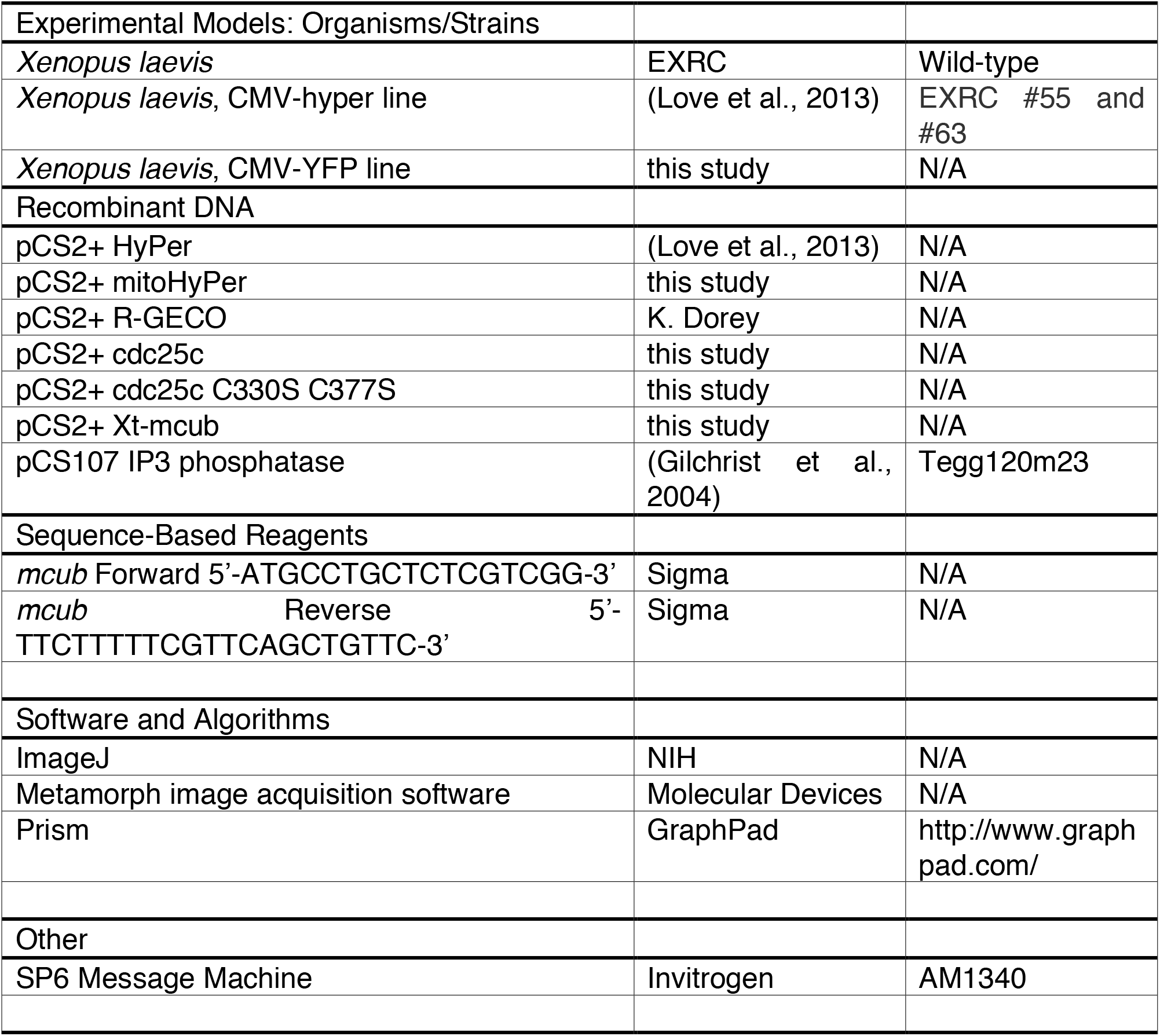

## EXPERIMENTAL MODEL AND SUBJECT DETAILS

All experiments involving animals were approved by the local ethical committee and the Home Office. Unfertilized oocytes were obtained from *X. laevis* wild type females by injecting with 250 units of HCG. Embryos were obtained from in vitro fertilization. Ovaries were isolated from females after they had been humanely sacrificed.

## METHODS

### Manipulation of *Xenopus* oocytes and embryos

RNA synthesis using SP6 Message Machine was carried out following the manufacturers protocol. Stage VI Oocytes were isolated manually from ovaries of PMSG-primed female frogs, injected with 20 ng HyPer or SypHer RNA, and cultured in OR2 medium at 16 °C for 24 hours. Then progesterone was added to the medium at a final concentration of 2 *μ*M for 16 hours at 16 °C. 20 ng HyPer RNA was coinjected with 20 ng IP3 phosphatase RNA or 20 ng MCUb RNA, cultured and matured as above. To inhibit MCU, oocytes were injected with 40 nl solution containing 0.4pM Ruthenium Red and 20 ng HyPer RNA, cultured in OR2 medium at 16 °C for 24 hours before in vitro maturation. Embryos were injected with 10 nl of 5 mM ruthenium red (50 pM) into one blastomere at 2-cell stage and cultured at 22°C in 0.1XMMR.

Oocytes injected with 100 pg human Cdc25c RNA for wild type or cysteine mutants were treated with progesterone immediate after injections. Matured oocytes were identified by presence of a white spot at the animal pole, associated with germinal vesicle breakdown. Oocytes were incubated with inhibitors for 20 min before prick activation.

The concentrations of inhibitors used here are based on previously published studies in tissue culture cells and determined in preliminary experiments.

### Plasmids construction

MitoHyPer and SypHer constructs were gifts from Dr. V. Belousov. The MitoHyPer ORF obtained by digesting with *Not*I followed by Klenow and *Nhe*I, and SypHer ORF obtained by digesting with *Hind*III followed by Klenow and *Nhe*I was subcloned into the pCS2+ vector. *X. tropicalis mcub (ccdc109b)* was cloned by RT-PCR. Human Cdc25C wild type and cysteine mutant obtained from Addgene (#10964 and #10965 respectively) and ORFs obtained by digesting with NcoI and XbaI were subcloned into pCS2+ vector.

### Ca2+ and HyPer Imaging

Oocytes were injected with 20 ng of R-GECO RNA, matured and imaged. Images were captured via a Nikon A1 microscope. The settings were as follows, pinholes [46 *μ*m], scan selection [Galvano], format [512 × 512]. Images DsRed were excited with the 543 nm laser lines. For laser wound, laser was set as follows: wavelength [561 nm], laser power [40], pulse [10]. Embryos from transgenic female expressing HyPer or oocytes injected with HyPer RNA were imaged using a Nikon TE2000 PFS microscope. Signals were excited with filters of BP430/24 and BP500/20, and detected with a filter of BP535/30 unless otherwise indicated. Images were analyzed by subtracting background, smoothing, formatting to 32-bit, then dividing using ImageJ. Since the HyPer ratio before activation vary among the different batches of oocytes, the ratio for control before activation was set as 1.0 and used for normalization for all samples.

### Detection of ROS by Amplex Red in oocytes

Albino oocytes were injected with 10 nl of solution containing 10 mM Amplex Red with or without 320 units/ml HRP. Time lapse movies were taken every 1 min after injection using Leica M205FA with DSR filter and the camera, DFC365FX.

### Measurement of ATP

ATP was measured using the ATP-determination Kit (Invitrogen) according to the manufacturer’s protocol. *Xenopus* laevis eggs were collected and immediately lysed with ice-cold 1X passive lysis buffer (Promega) and centrifuged at 13500 rpm for 10 min at 4 °C. A 10 *μ*L of the supernatant was diluted in 990 *μ*l water. Then 10 *μ*l of sample or 10 *μ*L ATP standard solution was added to 90 *μ*L of reaction buffer in each well of a 96-well plate. Luminescence was measured using Mithras LB 940 Multimode Microplate Reader. 1 unit/mL apyrase was employed in this assay as a negative control. All experiments were run in biological triplicates and technical duplicates, and the background luminescence was subtracted from the measurement. ATP concentrations in experimental samples were calculated from the ATP standard curve.

### Immunoblotting

6–10 embryos were frozen in dry ice, kept at −80 °C until use, and homogenized in the extraction buffer (100 mM NaCl, 50 mM Tris, 5 mM EDTA, 1% NP-40, pH 7.5) containing cOmplete Mini EDTA-free protease inhibitor (04693159001, Roche) and PhosSTOP phosphatase inhibitor (04906845001, Roche) and centrifuged at 13000 rpm for 1 min. The supernatant was dissolved in Laemmli’s sample buffer containing 5% b-mercaptoethanol and heated at 95°C for 5 min. Aliquots equivalent to 1−2 oocytes or embryos were subjected to 7.5% or 10% SDS-PAGE. Proteins on the gel were transferred to PVDF membrane and incubated with antibodies. The following antibodies were used for western blot analyses: *anti-Xenopus* Cdc25C mouse monoclonal (hybridoma, clone DG6 is gift from Dr J. Gannon, The Francis Click Institute, supernatant 1:500 dilution), anti-Xenopus Cyclin B2 mouse monoclonal (1:1000 dilution), anti-Phospho-cdc2 (Tyr15) (1:1000 dilution), anti-CDC2 (p34) Mouse monoclonal (1:1000 dilution), anti-Cdc25C (1:1000 dilution), anti-Cdc25C phospho S216 (1:1000 dilution), anti-a-tubulin (1:50000 dilution), anti-rabbit IgG/HRP (1:25000 dilution), and anti-mouse IgG/HRP (1: 25000 dilution). The signal was visualized using chemiluminescence (Immobilon, Millipore).

### Immunohistochemistry

Whole mount immunohistochemistry was carried out according to (Chalmers et al., 2003) with a modification in which pigmented embryos were bleached in 10% H_2_O_2_/PBS for 1 hour after fixation. The following antibodies were used: anti-Lamin B1 (1:500) with anti-rabbit Alexa Fluor 647 (1:500) and anti-a-Tubulin (1:2000) with anti-mouse TRITC (1:250). SYTOX Green was used to stain DNA (1:100). Cleared embryos with Murray’s were mounted inside of the ring made of vacuum grease (Dow Corning^®^ high-vacuum silicone grease, Sigma, Z273554) on a coverslip, and another coverslip was put on the top with a slight pressure. Confocal images were acquired by FV1000 (Olympus) using laser 488 and 543 with filter sets for Acridine Orange, TRITC and Alexa 647.

### Nuclear transplantation followed by HyPer/SypHer imaging

Oocytes injected with 20 ng HyPer or SypHer RNA were matured by adding progesterone and injected with sperm nuclei as described in (Amaya and Kroll, 1999). Successfully dividing embryos were imaged every 30 seconds for 3-4 hours using filter sets described above.

### Statistical Analysis

Statistical analyses were performed with Prism (GraphPad Software). Data were first checked for variance, and the appropriate statistical tests showed in each figure legend were used to generate *P* values. p values < 0.05 were considered significant; * p < 0.05, ** p < 0.01, *** p < 0.001, **** p < 0.0001.

## AUTHOR CONTRIBUTIONS

S.I. and E.A. conceived the study. Y.C and N.R.L. initially observed and imaged ROS generation in embryos. Y.H. designed and performed oocyte experiments and HyPer/calcium imaging with contributions from Y.C. and N.R.L. S.I. designed and carried out plasmids construction, AmplexRed assay, MCUb and cell cycle experiments, immunoblots and HyPer/SypHer imaging using nuclear transplantation. J.I.G. imaged ROS oscillation in embryos. S.I., N.R.L. and E.A. wrote the manuscript.

## ACKNOWLEDGMENTS

We thank Professor J.E. Ferrell, Jr. and Dr. J. Gannon for the *anti-Xenopus* Cdc25C antibody, and Dr. Peter March for his help with the microscopy. The Bioimaging Facility microscopes used in this study were purchased with grants from BBSRC, Wellcome Trust and the University of Manchester Strategic Fund. This work was supported by studentships and grant from The Healing Foundation (Y.H., N.R.L., Y.C., E.A.) and a Medical Research Council Research Project Grant (S.I., J.I.G., E.A.).

## SUPPLEMENTAL FIGURE LEGEND

**Figure S1.**
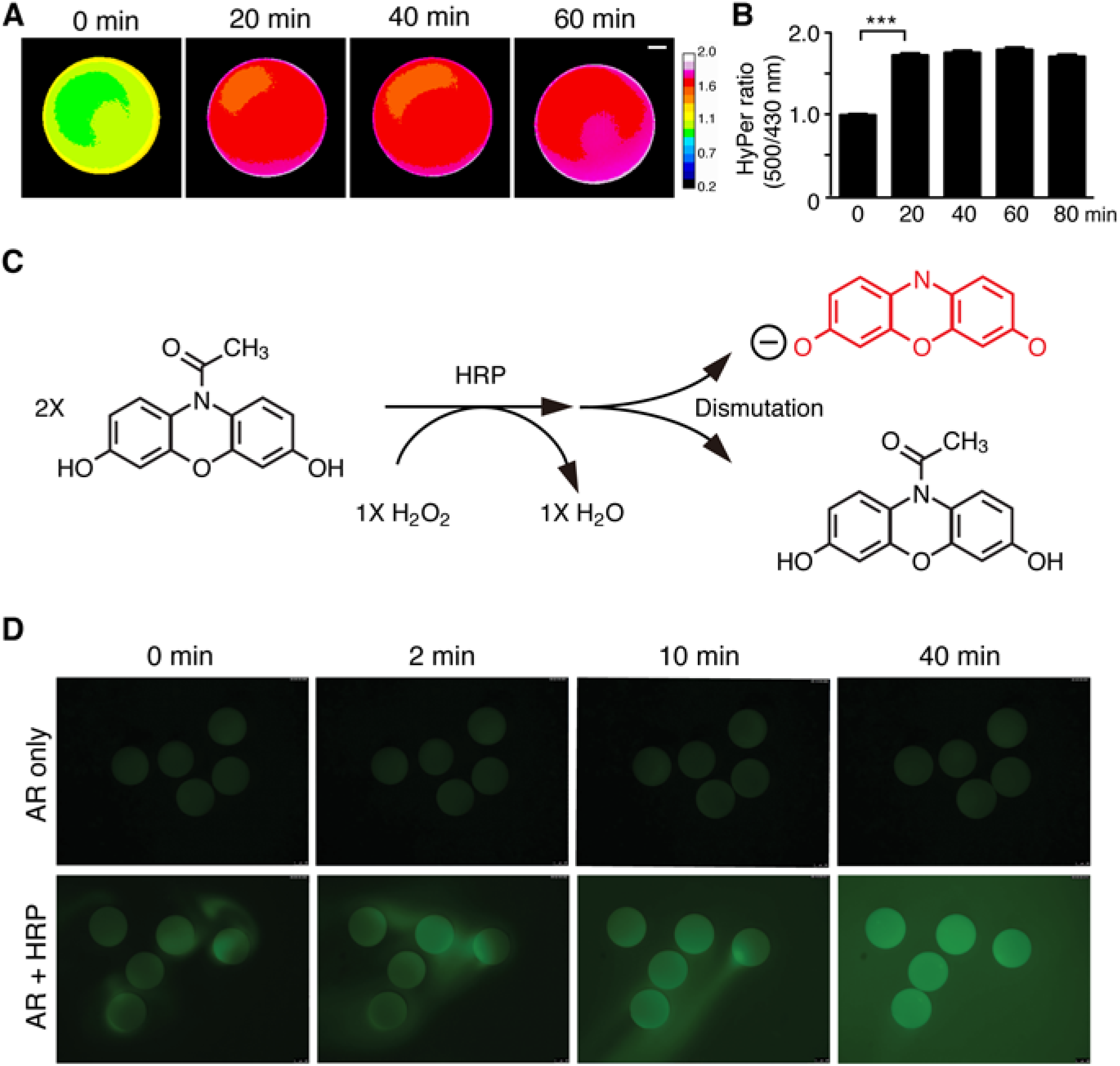
ROS are generated in oocytes after prick activation (related to Figure 1). (A) Oocytes obtained from a transgenic female expressing HyPer were prick-activated and imaged every 20 min. (B) Quantification of HyPer ratio in A. n = 60. ****p < 0.0001, 20 min compared to t0, paired t-test. Error bars represent mean ± SEM. (C) Amplex Red (AR) gives rise to fluorescent resorufin reacting specifically with H_2_O_2_ in the presence of horseradish peroxidase (HRP). (D) 10 nl of 10 mM AR was injected into albino mature oocytes with or without 320 units/ml HRP. The strong fluorescence (pseudo color) was observed just after AR injection with HRP. These pictures are from a representative of at least three independent experiments. See also Movie S2.

**Figure S2.**
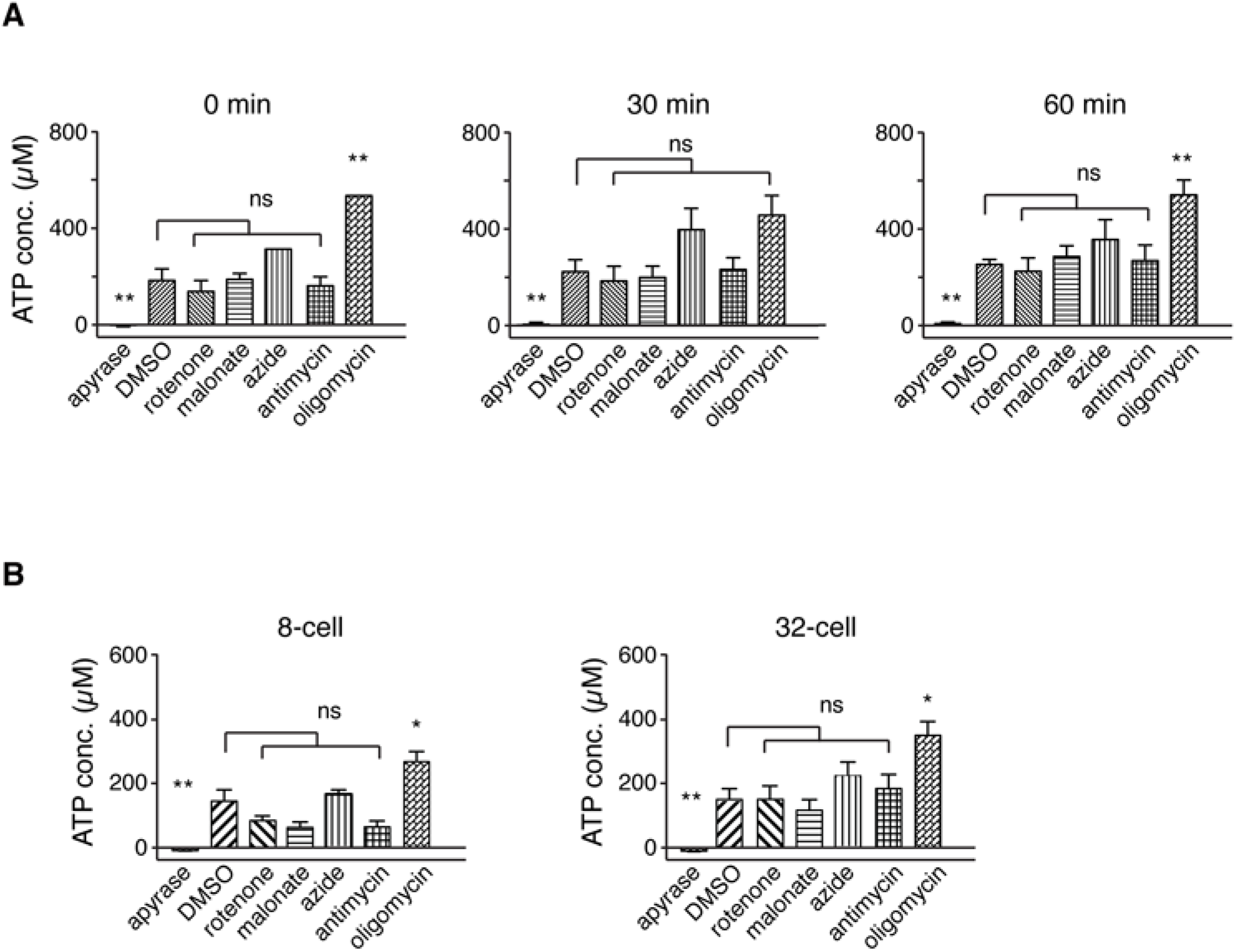
ATP levels are not affected by mtROS inhibitors (related to Figures 3 and 5). (A) The amount of ATP was measured at 0, 30 and 60 min after prick-activation in the oocytes treated with mitochondrial inhibitors. As a negative control, the extract was treated with apyrase ATPase. Three independent experiments were performed with technical duplicates. Error bars represent mean ± SEM. *p < 0.05, **p < 0.01, ****p < 0.0001; ns, not significant, compared to DMSO control, Unpaired t-test. For apyrase at 0 min and 30 min and azide at 60 min, Mann-Whitney test was used. (B) The amount of ATP in the embryos treated with mitochondrial inhibitors was measured at the 8-cell and 32-cell stages. There was no significant decrease of ATP by inhibitor treatment. Six independent experiments were performed with technical duplicates. Error bars represent mean ± SEM. *p < 0.05, **p < 0.01; ns, not significant, compared to DMSO control, Unpaired t-test. For apyrase, Mann-Whitney test was used.

**Figure S3.**
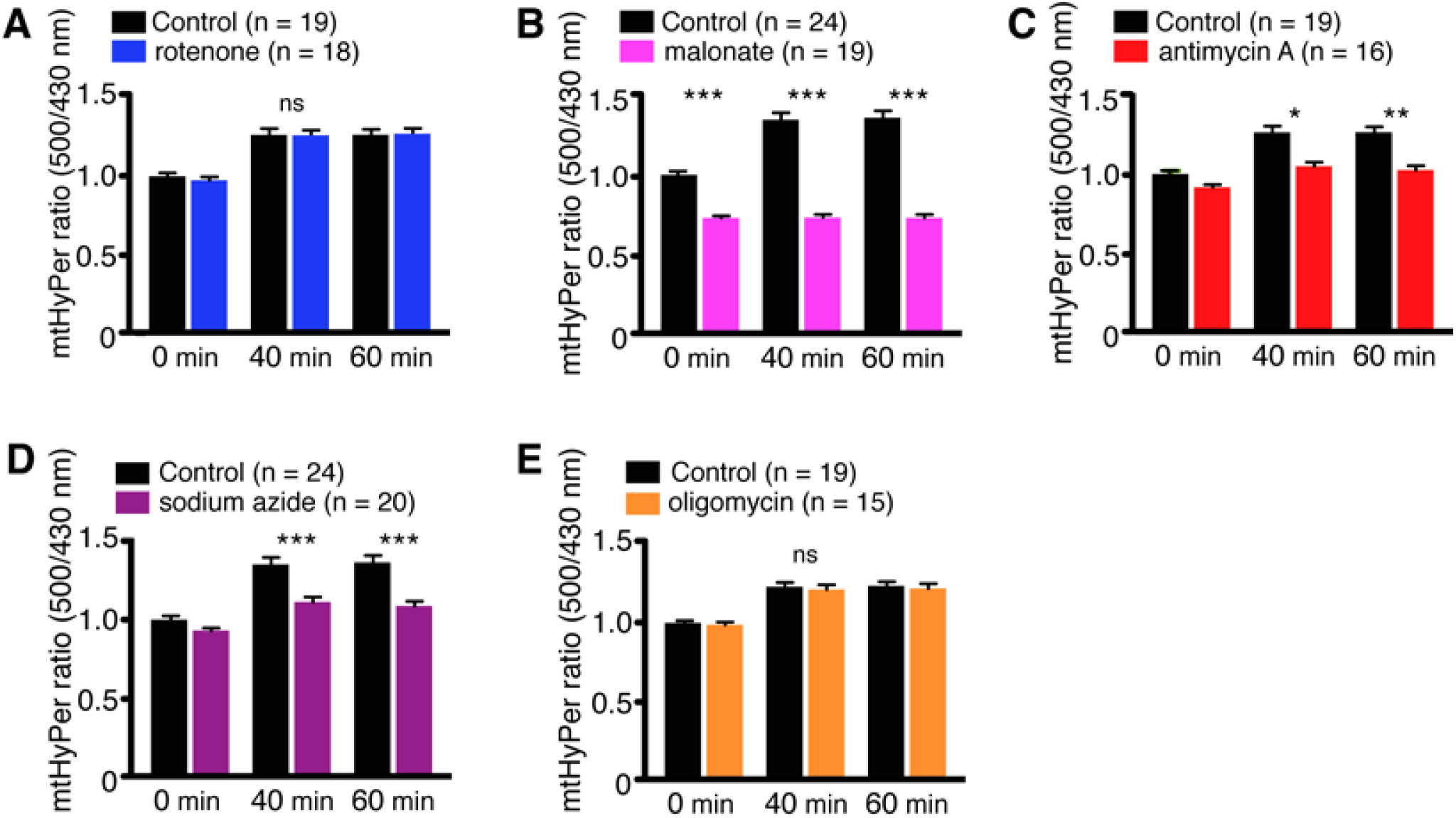
SypHer ratio in mature oocytes treated with mitochondrial inhibitors (A-E) and decreased ROS production detected by mitoHyPer in mature oocytes treated with mitochondrial inhibitors (F-J) (related to Figure 3). (A-E) Quantification of normalized SypHer ratio of oocytes treated with 1 *μ*M rotenone (A), 10 mM malonate (B), 10 *μ*M antimycin A (C), 3 mM sodium azide (D) and 6 *μ*M oligomycin. No significant change of SypHer ratio was observed (two-way ANOVA). (F-J) Quantification of normalized mitoHyPer ratio of oocytes treated with 1 *μ*M rotenone, no significant decrease in ratio was found (A), 5 mM malonate, 19.8% at 0 min, 34.6% at 40 min and 35.7% at 60 min decrease in ratio (B), 10 *μ*M antimycin A, 14.5% at 40 min and 16.5% at 60 min decrease in ratio (C), 1 mM sodium azide, 13.1% at 40 min and 14.9% at 60 min decrease in ratio (D) and 6 *μ*M oligomycin, no significant decrease in ratio was found (E). Error bars represent mean ± SEM. ****p < 0.0001; ns, not significant. two-way ANOVA and Sidak post hoc tests.

Movie S1. Time-lapse movie showing ROS production from fertilization to the blastula stages in transgenic embryo expressing HyPer (related to Figure 1). Signal was obtained using excitation filters of 490/20 nm (left) and 402/15 nm (middle) with an emission filter of BP525/36. The HyPer ratio was calculated using ImageJ (right).

Movie S2. ROS detection using Amplex Red (AR) (related to Figure 1). Albino oocytes were injected with AR only (left) and AR with HRP, and time lapse movies were taken every 1 min after injection using Leica M205FA with DSR filter (pseudo color).

Movie S3. Detection of Ca^2+^ wave with R-GECO after laser activation. Image of immature (left) and mature (mature) oocytes injected with mCherry (3 seconds of movie) or R-GECO (3 seconds of movie) (related to Figure 2). Ca^2+^ wave was observed in the R-GECO movie, but only in the mature oocyte (right).

Movie S4. Ca^2+^ wave is induced by addition of Ca^2+^ ionophore, A23187 in mature oocyte (right). Ethanol alone control does not induce Ca^2+^ wave (left). (Related to Figure 2).

Movie S5. Ca^2+^ wave induced by laser in mature oocyte is disrupted by the presence of EGTA. (Related to Figure 2).

Movie S6. Overexpression of IP3 phosphatase in oocyte impairs Ca^2+^ wave. (Related to Figure 2).

Movie S7. Mitochondrial inhibitors do not disrupt Ca^2+^ wave. (Related to Figure 3).

Movie S8. Overexpression of MCUb RNA does not inhibit Ca^2+^ wave. (Related to Figure 4).

Movie S9. Removal of inhibitors, 10 mM malonate and 3 mM sodium azide restores the ROS production in HyPer transgenic embryos. Time lapse movies were taken every 30 sec after 4-cell arrested embryos were transferred to medium without inhibitors. (Related to Figure 6).

Movie S10. Oscillation of ROS along with cell division. Sperm solution was added to unfertilized oocyte expressing HyPer, and imaged every 30 sec for 5 hours. Images were processed without smooth using ImageJ and Brightness/Contrast was set between 1.2 and 1.6. (Related to Figure 7F).

Movie S11. Another example of oscillation of ROS in embryo expressing HyPer. A dividing embryo after fertilization was imaged every 30 sec for 5 hours with 1000 ms exposure for YFP500 and 500 ms for CFP430. Images were processed without smooth and Brightness/Contrast was set between 2.9 and 3.8. (Related to Figure 7F).

